# Environmental and biotic drivers of *Aedes albopictus* spatiotemporal distribution in the Argentina-Brasil-Paraguay subtropical triple border: The key role of periurban and disturbed wild environments

**DOI:** 10.64898/2026.02.02.703241

**Authors:** Julieta Siches, M. Victoria Micieli, Pablo E. Berrozpe, Maria del Rosario Iglesias, Juan J. García, María V. Cardo

## Abstract

There is empirical evidence that biophysical factors determine the spatio-temporal distribution of mosquito vectors, and identifying the variables that shape their ecology allows decision-makers to design effective surveillance and control strategies. This study evaluated the spatiotemporal distribution of *Aedes albopictus* in relation to environmental and biotic variables in the Iguazú Department, Misiones Province, Argentina, within the tri-border region shared with Brazil and Paraguay. Environmental characterization integrated field data and remotely sensed biophysical variables, and vector occurrence was analyzed at micro- and meso-spatial scales using generalized linear mixed models. Eleven sampling sessions were conducted between April 2019 and February 2020 at 81 sites representing urban, periurban, and wild environments. A total of 1,614 *Ae. albopictus* and 4,358 *Ae. aegypti* specimens were identified. Rainfall, minimum temperature, exposure days, and land cover were the main predictors of *Ae. albopictus* presence, showing nonlinear responses to precipitation and vegetation. The selected model explained 67% of the variance. The species exhibited clear spatiotemporal stratification, with periurban and disturbed wild areas functioning as ecotones favorable to its establishment. These findings provide key insights to guide preventive actions and strengthen integrated vector management strategies in the region.

## Introduction

Over the past five decades, arboviral diseases —such as dengue, Zika, yellow fever and chikungunya— have emerged as some of the most widespread and pressing public health challenges in the Americas. These infections have been particularly burdensome in tropical and subtropical regions, where they account for the highest rates of morbidity and mortality [1]. Transmission of these vector-borne diseases depends on the simultaneous presence of the host, the vector, and the pathogen within the same geographic area [2]. Moreover, transmission dynamics are influenced by the diversity, abundance, and spatial distribution of both hosts and vectors within the landscape, resulting in micro-spatial heterogeneity in transmission patterns [3]. Understanding the ecological characteristics of disease vectors is therefore crucial for identifying their distribution patterns and for developing more effective strategies to address current and emerging epidemiological threats.

*Aedes albopictus* (Diptera: Culicidae, Stegomyia) is an invasive species native to Southeast Asia, recognized as a competent vector for a wide range of arboviruses. These include the four dengue virus serotypes as well as viruses responsible for Japanese encephalitis, Potosi, Keystone, Tensaw, eastern equine encephalitis, yellow fever, chikungunya and Zika (4, 5). Notably, the potential role of *Ae. albopictus* in chikungunya virus transmission has gained particular attention in the American continent, where it has been shown to be a more effective vector than *Ae. aegypti* [6]. This is specially concerning in regions where its geographic range has recently expanded, such as parts of Europe [7].

The most likely pathway for the introduction of *Ae. albopictus* into the Americas is through the international trade of used vehicle tires with Southeast Asia. In 1985, the species was first recorded in Harris County, Texas, United States [8]. Within just two years, it had been reported in 15 of the 50 US states and by 1997, its distribution had extended to 25 states [9]. In South America, the first report occurred in Brazil in 1986 [10] and, by 2002, it had been detected in 20 of the 27 Brazilian states [11].

In Argentina, *Ae. albopictus* was first recorded in 1998 in San Antonio, Misiones Province, located in the northeast of the country [12]. Since then, its presence has been confirmed in a few additional localities within Misiones Province and a small area of neighboring Corrientes Province, with the current distribution extending between latitudes 25° 28° and 28° 10°S. In other words, twenty-five years after its first detection in Argentina, the distribution of *Ae. albopictus* remains limited, with only a few locations showing signs of stabilization [13, 14]. Although several studies have explored potential factors restricting its distribution, including competitive exclusion, the composition of container-breeding mosquito communities, environmental preferences, and egg diapause [13, 15], the reasons behind the failure of *Ae. albopictus* to expand in Argentina still remain uncertain.

Health risks linked to biodiversity loss and ecosystem degradation have been repeatedly highlighted, such as the increased burden of vector-borne diseases resulting from land degradation and land cover transformation [16]. For instance, in South America, land-use changes such as road construction have been strongly associated with increased incidence of malaria, leishmaniasis, and other parasitic diseases among workers [17]. Development projects can increase human exposure to natural mosquito habitats. On the other hand, they can also create more suitable habitats for the proliferation of these insects, such as forest edges and associated microclimates [18]. The conversion of wild forests into agricultural land, in particular, has been shown to exacerbate the spread of mosquito-borne diseases [17].

Numerous studies suggest that *Ae. albopictus* population densities fluctuate interannually in response to climatic variability [19]. For example, the highest abundances have been observed at temperatures around 25 °C (20), while conditions simulating unfavorable winter scenarios such as short photoperiods (8 h light: 16 h dark) and temperatures of approximately 16 ± 2.3 °C have been shown to inhibit completion of its life cycle [21]. Adult *Ae. albopictus* abundance has also been positively correlated with the density of larval habitats, which is in turn influenced by microclimatic conditions [20]. Precipitation plays a key role by increasing the number of outdoor containers that accumulate rainwater, thereby creating suitable oviposition sites. Vegetation indices are frequently used in mosquito studies; specifically, the Normalized Difference Vegetation Index (NDVI) has been consistently associated with infestation levels of both immature and adult *Ae. albopictus* [15, 19]. *Ae. albopictus* abundance has also been reported to be strongly associated with socio-ecological factors, including the accessibility and quality of public services, vegetation degradation, and the state of urban infrastructure [22].

For its part, *Ae. aegypti* (L.) is widely recognized as the primary vector of several major arboviral diseases in urban environments, both in the Americas and globally [23]. In Argentina, its distribution is broad and expanding, encompassing the entire northern and central regions of the country as well as reaching the northern limits of the Patagonian region [24]. Coexistence between *Ae. albopictus* and *Ae. aegypti* in the same larval containers has been reported [13], and in some contexts, *Ae. albopictus* has been hypothesized to competitively displace *Ae. aegypti* [25]. However, such displacement has not been observed in Argentina, where recent studies conversely indicate that *Ae. aegypti* may outcompete *Ae. albopictus* in urban environments or that the two species may exhibit co-dominance in rural settings [13]. Habitat segregation has been proposed as a mechanism facilitating coexistence by reducing direct competition between species [26]. In other regions, *Ae. aegypti* predominates in urban environments, whereas *Ae. albopictus* is more common in rural areas, with both species overlapping in periurban zones [27]. This segregation pattern could be attributed to a preference of *Ae. albopictus* for environments with certain characteristics, such as the availability of particular breeding sites, a greater diversity of food sources, and a higher number of shelters, rather than solely by competitive exclusion. This is especially relevant considering that *Ae. albopictus* exhibits a more peridomiciliary behavior, in contrast to the intradomiciliary preference of *Ae. aegypti* [28].

The study area, Puerto Iguazú, is located in the northwesternmost part of Misiones Province, Argentina, at the tri-border area shared with Brazil and Paraguay. During recent years, the city has experienced rapid urban expansion characterized by uneven, disordered, and unplanned growth, resulting in limited access to essential public services such as potable water, sewage, electricity, and waste management. Iguazú National Park, located 17 km apart from Puerto Iguazú, is one of the most biodiverse protected areas in Argentina and boasts the highest number of endemic species in the country [29]. Despite its protected status, the park is impacted by human activity, including tourism infrastructure and housing for park staff. Managing and preserving this natural space poses major challenges, especially given the high volume of visitors [29]. These challenges include increased human–wildlife interactions, which elevates the risk of exposure to both known and emerging zoonotic diseases and management waste [30].

From an epidemiological perspective, the Iguazú Department holds strategic importance due to its high volume of human movement across land and river borders, as well as through its airport, which operates both domestic and international flights. The latter has been identified as a major contributor to the spread and intensification of arboviral diseases such as dengue [31]. Cases of both imported and autochthonous dengue have been consistently reported from the 2015/16 to the 2023/24 seasons, with the exception of the 2016/17 season [32]. Additionally, the 2022/23 season recorded a notable outbreak of chikungunya fever [33]. This underscores the Iguazú Department’s critical role as both a point of entry and establishment for circulating arboviruses and highlights the potential role of *Ae. albopictus* as a bridge vector between sylvatic transmission cycles and periurban/urban environments [34, 35].

In this context, gaining insights into the ecology of disease vectors is essential for deciphering their distribution patterns and developing more effective strategies to address emerging epidemiological challenges. The objective of the present study was to assess the environmental and biotic factors associated with the spatiotemporal distribution of *Ae. albopictus* in northeastern Argentina, using the city of Puerto Iguazú and the Iguazú National Park as a study case. This study is guided by four main lines of inquiry, concerning the association between *Ae. albopictus* distribution and the degree of anthropogenic intervention, climatic factors, land cover class, and the presence of *Ae. aegypti*. Addressing these lines, will contribute to a deeper understanding of the determinants shaping the distribution of *Ae. albopictus*, and clarify the constraints on its expansion, with the aim to propose vector control and prevention tools for mitigating zoonoses transmission. This knowledge is particularly relevant in the context of recent climatic changes, intensified by global warming, which pose profound implications for the health of both human populations and ecosystems.

## Materials and methods

### Study area

Puerto Iguazú is located in the Paranaense Forest ecoregion, which is part of the Atlantic Forest complex in Misiones Province, Argentina (Fig. 1). This ecoregion is characterized by a semi-deciduous forest with differentiated tree strata, abundant epiphytes, bamboo, and lianas. The climate is subtropical, with an average annual temperature of 21 °C, ranging from 24 °C in summer to 14 °C in winter. Annual precipitation is approximately 1.800 mm, concentrated in the summer season [36].

**Figure 1.**
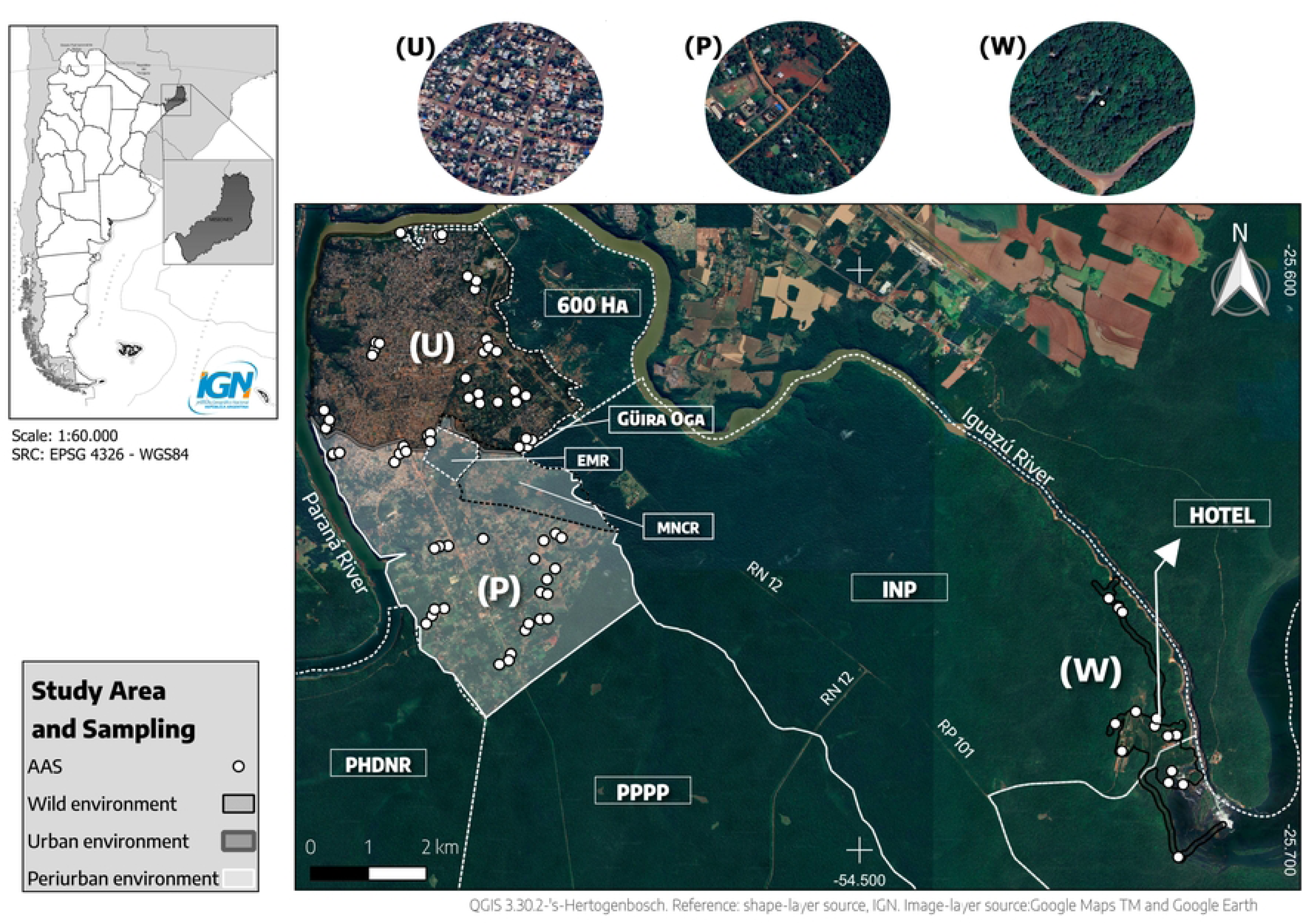
Study area. categorized a priori in three environmental types, urban (U), periurban (P) and wild (W), and surrounding natural areas, including: Puerto Peninsula Provincial Park (PPPP), the Peninsula Harbor Defense Nature Reserve (PHDNR), Iguazu National Park and Reserve (INP), Mboreré Municipal Natural and Cultural Reserve (MNCR), “El Eucaliptal” Municipal Natural Reserve (EMR), Andrés Giai Protected Landscape (Güira Oga), and the Iriapú Reserve (600 Ha). The primary routes and roads in the study area are RN 12 and RP 101. The distribution of adult activity sensors (AAS) is also indicated on the map (white dots).

### Entomology sampling

Based on the visual analysis of high-resolution satellite imagery and national sociodemographic data obtained from the official statistics and census institute [37, 38, 39], for sampling purposes the study area was classified a priori into three environmental types: urban (U), characterized by high population density and infrastructure, corresponding to the central area of Puerto Iguazú; periurban (P), defined by a mix of rural and urban elements with lower population density and subsistence agricultural activities; and wild (W), featuring predominant native vegetation and minimal anthropogenic intervention, encompassing the public use and trail areas of the Iguazú National Park (INP). A vector layer of polygons, composed of geometric shapes used to represent spatial features, was created on the image to delineate these environments (Fig. 1). This classification was further evaluated by ordering the sampling units in a multivariate space as a function of the environmental variables recorded in a Principal Component Analysis for a posteriori classification (see below).

Sampling was conducted from April 2019 to February 2020 on a monthly basis, obtaining samples for all seasons of the year. Ovitraps consisting of 1 L black plastic containers lined with corrugated cardboard to support egg deposition were employed as adult activity sensors (AAS). The study area was divided into 400 m² quadrants, which were randomly selected to cover 10% of the total area in each of the three defined environmental types. Four AAS were randomly placed in each selected quadrant, with the exception of quadrants U3, P7, P3 and W4, where three AAS were deployed, and W5, which had only one AAS, due to accessibility issues. Therefore, a total of 35, 30, and 16 AAS were deployed in U, P, and W, respectively. They were positioned at 1 -1.5 m height [40], filled to 50% of their capacity with dechlorinated tap water, and left uncovered for 6 to 8 days to allow oviposition. After this period, the AAS were covered (deactivated) until the next sampling cycle. Any missing or damaged AAS were replaced with new ones. After the last sampling event, the traps were dismantled.

The egg supports were transported to the insectary at the National Tropical medicine institute (INMeT, ANLIS Malbrán - MSAL) in sealed, properly labeled bags. Upon arrival, the supports were spread out and allowed to dry in an incubator at 19 ± 2 °C until immersion. The hatched larvae were reared in 250 ml plastic containers covered with semi-transparent polyester mesh that permitted air circulation and light exposure and fed a standard laboratory diet. Once the larvae reached the pupal stage, they were transferred to smaller containers (20 ml capacity), which were placed in emergence cages. The emerged adults were euthanized with acetone in a lethal chamber, dried, mounted using ad-hoc techniques, and identified under a stereoscopic microscope using dichotomous keys [41].

### Explanatory variables

#### Macrohabitat

Spectral indices were used to characterize the land surface based on its spectral response, providing information on vegetation greenness, moisture, photosynthetic activity, and the presence of built-up areas or bare soil. For each season -defined as Autumn (April-June), Winter (July-September), Spring (October-December), and Summer (January-February)- a preliminary processing stage was conducted using the Google Earth Engine platform. This consisted of filtering Sentinel-2 Level 2A scenes (10 m spatial resolution) based on the area of interest, the sampling period, and a threshold for minimal cloud cover. The NDVI, which assesses vegetation health [42]; the Normalized Difference Water Index (NDWI), designed to delineate and enhance the presence of open water bodies [43]; ad the Coloration Index or Saturation Index (CI), developed to identify areas of bare soil and assess their degradation [44], were calculated. Additionally, a land surface temperature (LST) layer for the area of interest was constructed using Landsat 8-OLI (Level 2) imagery, specifically from the TIR 1 band (B10) with a spatial resolution of 100 m. These layers were imported into QGIS 3.16.4, where data were extracted based on the vector point layer corresponding to the georeferenced locations of the deployed AAS.

To create high-resolution land cover maps, PlanetScope imagery with a 3 m spatial resolution was used [45]. Supervised classifications were performed using QGIS 3.16.4 with the Semi-Automatic Classification Plugin (SCP). The supervised classification method employed was the Minimum Distance algorithm, utilizing the NDVI index along with the green, blue, red, and infrared bands. Four classes were identified: Impervious, low vegetation (LowVeg), high vegetation (HgVeg) and bare soil (Soil). The resulting land cover map was clipped in a 150 m radius buffer around each AAS, and the percentage area of each land cover type per sampling site was calculated using the QGIS zonal histogram algorithm. Also, a vector layer was manually drawn based on Google Earth images to identify water bodies within the study area, and the distance from each AAS to the nearest water body was calculated (Supplementary data 1).

Precipitation data was considered at two time lags, 14 and 30 days prior to AAS activation, based on records from the Puerto Iguazú airport weather station [46]. Selected demographic variables were obtained from National Institute of Statistics and Censuses [39] dataset, as detailed in Table 1. Elevation was not included as a variable in the analysis, as it was not considered a significant factor in the study area.

**Table 1.** Enviromental variables.

| *Variable* | *Description* | *Source* | *Unit* | *Used in* |
| --- | --- | --- | --- | --- |
| ***Climatic*** |  |  |  |  |
| *rainfall_30* | Precipitation accumulated 30 days before the closure of each monitoring period | SMN | mm | GLMM |
| *globtemp* | Average air temperature | in situ | °C | GLMM |
| *globhum* | Average percentage of relative humidity | in situ | % | GLMM |
| *globwind* | Average wind speed | in situ | m/s | GLMM |
| *globlight* | Average light intensity | in situ | klux | GLMM |
| *Min_T* | Average minimum temperature recorded during the week of monitoring | SMN | °C | GLMM |
| *Max_T* | Average maximum temperature recorded during the week of monitoring | SMN | °C | GLMM |
| ***Environmental*** |  |  |  |  |
| *watertemp* | Water temperature | in situ | °C | GLMM |
| *oxy* | Dissolved oxygen in the water | in situ | OD | GLMM |
| *pH* | Water pH | in situ |  | GLMM |
| *sal* | Salinity/conductivity | in situ | ppt | GLMM |
| ***Vectorial*** |  |  |  |  |
| *albobin* | Presence/absence of *Aedes albopictus* | in situ | - | GLMM |
| *aegbin* | Presence/absence of *Aedes aegypti* | in situ | - | GLMM |
| *aeg_day* | Number of *Ae. aegypti* specimens collected, standardized by sensor active days. | in situ | - | GLMM |
| ***Geographical*** |  |  |  |  |
| *Road_type* | Road types by category: paved, dirt, cobblestone and gateway | in situ | - | GLMM |
| *environment* | Environment by category: urban, periurban, wild environment and disturbed wild environment | by authors | - | GLMM |
| *gateway* | Road type where the AAS is located: Gateway | in situ | - | PCA |
| *paved* | Road type where the AAS is located: Paved Road | in situ | - | PCA |
| *cobblestone* | Road type where the AAS is located: Cobblestone | in situ | - | PCA |
| *dirt* | Road type where the AAS is located: Dirt Road | in situ | - | PCA |
| *NDVI* | Green Normalized Difference Vegetation Index | Sentinel 2 (level 2A) + GEE | - | GLMM |
| *NDWI* | Green Normalized Difference Water Index | Sentinel 2 (level 2A) + GEE | - | GLMM |
| *LST* | Land Surface Temperature | Landsat 8-OLI (Level 2) + GEE | °C | GLMM |
| *NDVI_win* | Normalized Difference Vegetation Index for winter | Sentinel 2 (level 2A) + GEE | - | PCA |
| *NDWI_win* | Normalized Difference Water Index for winter | Sentinel 2 (level 2A) + GEE | - | PCA |
| *CI_win* | Coloration Index for winter | Sentinel 2 (level 2A) + GEE | - | PCA 31 |
| *NDVI_sum* | Normalized Difference Vegetation Index for summer | Sentinel 2 (level 2A) + GEE | - | PCA |
| *NDWI_sum* | Normalized Difference Water Index for summer | Sentinel 2 (level 2A) + GEE | - | PCA |
| *CI_sum* | Coloration Index for summer | Sentinel 2 (level 2A) + GEE | - | PCA |
| *Water_m* | Distance to the nearest water body | Planet Scope 3 m + QGIS | m | GLMM |
| *Impervious* | Impervious class | Planet Scope 3 m + QGIS | m<sup>2</sup> | GLMM & PCA |
| *LowVeg* | Low vegetation class (includes herbaceous or low shrubs, typically <1.5 m height) | Planet Scope 3 m + QGIS | m<sup>2</sup> | GLMM & PCA |
| *HgVeg* | High vegetation class (includes tall shrubs and trees, typically >1.5 m height) | Planet Scope 3 m + QGIS | m<sup>2</sup> | GLMM & PCA |
| *Soil* | Bare soil class | Planet Scope 3 m + QGIS | m<sup>2</sup> | GLMM |
| **Demographic** |  |  |  |  |
| *totalpop* | Total population | INDEC 2010 | inhab | GLMM & PCA |
| *unoccupied_rate* | Unoccupied rate | INDEC 2010 | - | GLMM & PCA |
| *totalmale* | Total male population | INDEC 2010 | inhab | PCA |
| *totalfemale* | Total female population | INDEC 2010 | inhab | PCA |
| *households* | Total number of households | INDEC 2010 | - | PCA |
| *privdwellings* | Total number of private dwellings | INDEC 2010 | - | PCA |
| *o_priv_dwell* | Total number of occupied private dwellings | INDEC 2010 | - | PCA |
| *house* | Total number of houses | INDEC 2010 | - | PCA |
| *rancho* | Number of dwelling types: 'Rancho' (houses with adobe walls, earthen floors, and tin or thatch roofs). (INDEC - SESD Glossary, 2010) | INDEC 2010 | - | PCA |
| *casilla* | Number of dwelling types: 'Casilla' (small houses built with low-quality or scrap materials). (INDEC - SESD Glossary, 2010) | INDEC 2010 | - | PCA |
| *apartment* | Dwelling type: Apartment | INDEC 2010 | - | PCA |
| *prem_not_built_dwell* | Dwelling type: Premises not built for dwelling | INDEC 2010 | - | PCA |
| *room_tenants* | Dwelling type: Room with tenant | INDEC 2010 | - | PCA |
| *motorhome* | Dwelling type: Motorhome | INDEC 2010 | - | PCA |
| **Time** |  |  |  |  |
| *days* | Number of days that the AAS was active | Time | Date | GLMM |
| *month* | Monitoring months | Time | Date | GLMM |

a. Environmental variables used to characterize the sites where the adult activity sensors (AAS) were placed, both for the assessment of site classification by principal component analysis (PCA) and for the generalized linear mixed modeling (GLMM) of *Ae. albopictus* occurrence in Puerto Iguazú, Argentina. For climatic variables measured in situ, averages were calculated from values recorded at the beginning and end of sensor activity. For variables obtained from INDEC, values were taken from the calculations for census tracts (which are geo-statistical units defined by the organization for the national census).

#### Microhabitat

Upon activation of each AAS, the structural characteristics of the placement site (asphalt, cobblestone, dirt, or walkway) were documented, and the following parameters were recorded using a thermohygrometer: air temperature (°C), relative humidity, wind speed (Anemometer Peakmeter - Pm6252a®), and light intensity (Luxometer CEM DT-8809A®). At the end of the active period, the previous measures were repeated along with physicochemical properties of the water, including pH, dissolved oxygen (DO), salinity, and water temperature (Multiparameter probe AQUACOMBO HM3070®) (Table 1).

### Statistical analyses

All analyses were performed using the free software platform R [47].

To evaluate the classification of the three environmental types conducted prior to sampling, a Principal Component Analysis (PCA) was performed using the package stats [47]. In this analysis, “sites” (rows) corresponded to each site where the AAS were placed, while “variables” (columns) included environmental factors such as demographic data, infrastructure details (e.g., road types), land cover and land use types, and proximity to water bodies, as outlined in Table 1.

The infestation status of *Ae. albopictus* and *Ae. aegypti*, both overall, monthly and per environmental type, was assessed by the Positivity Ovitrap Index (POI, Formula 1), commonly used in epidemiology to gauge the risk of arboviral transmission [48]. Three POI ranges were defined as indicators of low (POI ≤ 40%), moderate (41-60%) and high (>60%) entomological risk.

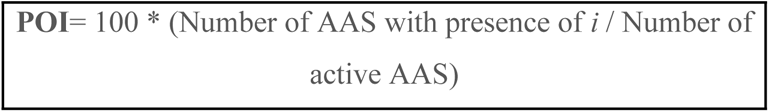

With *i* being either *Ae. albopictus* or *Ae. aegypti*. Presence: at least 1 egg or 1 larva

To assess the co-occurrence of *Ae. albopictus* and *Ae. aegypti* per environment, the significance of their association was evaluated using an adapted version of the Bayesian Proportion Difference Test proposed by Risso & Risso (2017) [49]. The null hypothesis (H0) was defined as Co-Occurrences (CO, indicating that *Ae. aegypti* and *Ae. Albopictus* are both present), Exclusive Occurrence (EO, indicating that *Ae. albopictus* is present given that *Ae. aegypti* is absent) and implies no association between species, while the alternative hypothesis (H1) was defined as CO > EO, indicating association. Subsequently, for environments presenting association (p<0.05) a diagnostic validation test was conducted using the Odds method. This methodology assesses the presence or absence of an “event” (herein, the occurrence of *Ae. albopictus*) based on a dichotomous “effect” (the occurrence of *Ae. aegypti*) and was performed at the monitoring event scale (AAS x month). The posterior odds were calculated, indicating the likelihood of *Ae. albopictus* being present in the absence/presence of *Ae. aegypti*. The Probability of Co-Occurrence (PCO), which in this context is the probability of both species being present, was then compared with the EO to ascertain the direction of the association. If PCO > EO, it indicates co-existence; otherwise, it indicates avoidance.

### Generalized Linear Mixed Models

The occurrence of *Ae. albopictus* was analyzed using generalized linear mixed models (GLMM) with a binomial error distribution using the MASS package [50]. Initially, a pairwise correlation analysis was performed among all explanatory variables considered for inclusion in the model, using a cutoff of *ρ* = 0.6. Subsequently, univariate models were run to assess the strength of the association between each explanatory variable and the response variable. The model was built using stepwise selection, with explanatory variables (if continuous, previously standardized) introduced one by one, along with their interactions and squared terms for continuous variables. The fixed effects evaluated included the NDVI, land cover classifications for each AAS, distance to water bodies, presence and abundance of *Ae. aegypti* (standardized by the number of days that the AAS was active), the rate of unemployment and total human population, precipitation, and environmental variables recorded *in situ* (Table 1). Random effects considered were each AAS (labelled ID), the sampling quadrants, and the four seasons. Model selection was based on the Akaike Information Criterion (AIC), with a delta AIC >2 considered significant [51]. To assess multicollinearity among explanatory variables, the variance inflation factor (VIF) was calculated at each modeling step, and variables were only retained in the model if their VIF values were below 5 [52]. The residuals of the selected models were visually inspected to check that they follow model assumptions. Concordance was evaluated using Cohen’s Kappa estimators [53]. To measure the proportion of variability explained by fixed and random factors, conditional and marginal R² values were calculated, representing the variance explained by fixed effects and by the full model (fixed + random), respectively, using the MuMIn package [54]. Finally, a spatial interpolation was performed through a spatial autocorrelation analysis using the POI values by ID and season. Since sampling points located in the INP were far apart from the urban nucleus, autocorrelation was assessed separately in INP points and non-INP points by computing Moran’s I correlograms. If significant autocorrelation was detected, a theoretical semivariogram was fitted and subsequently used to perform interpolation through the Kriging method, implemented with the gstat package [55]. If no spatial autocorrelation was found, interpolation was carried out using the Inverse Distance Weighting (IDW) method, also available in gstat.

## Results

### Environmental types. Validation and reclassification

The cumulative variance percentage of the first three principal components (PC) of the PCA was 70.1% (PC1=38.4%, PC2=18.7%, PC3=13.0%). The sites where the AAS were deployed were clustered into four main groups, based on the first two PCs. The group hereafter named “urban” included 26 out of the 35 sites previously classified as U and four sites previously classified as P (Fig. 2). The cluster was defined by demographic variables such as housing types, unemployment rate, number of households and residences, and population density, showing high correlation among them as well as land cover classification predominantly impervious and bare soil, with high NDWI values and, to a lesser extent, types of paved and asphalt roads. Sites hereafter named “periurban” were 26 previously classified as P and eight previously U sites (Fig.1). These 34 sites were grouped based on intermediate vegetation cover, lower population density, higher unemployment rate, and more precarious housing types, with less emphasis on unpaved and cobblestone roads. The a priori defined W category included six sites resembling P areas, located at the entrance of the INP along the main paved road and near the hotel (Fig. 1). These sites, characterized by high pedestrian traffic and people concentration, are hereafter referred to as “disturbed wild”, along with one site previously identified as U located on the border of Güira Oga and MNCR (Fig. 1). The subset of the remaining 10 W sites, hereafter “pristine wild”, were characterized by the presence of high vegetation, high NDVI values, and footpath roads.

**Figure 2.**
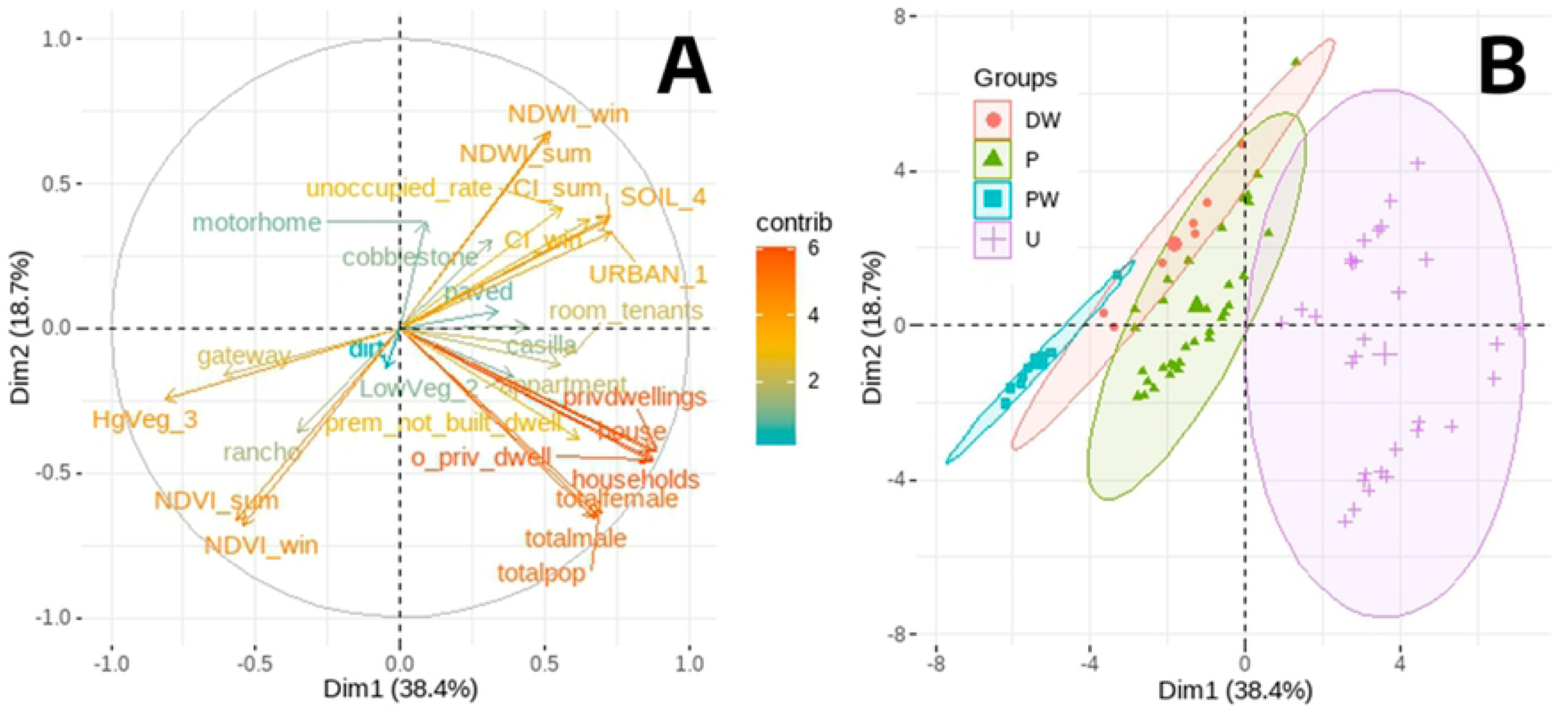
Biplot of the principal component analysis displaying the ordination of AAS deployment sites (B) as a function of environmental variables (A). The direction and length of the vectors indicate the magnitude of each variable’s contribution to the first two principal components, PC1 and PC2. The percentage of the total variance explained by each PC is indicated next to each axis. In (B), points indicate AAS deployment sites classified by environmental type: disturbed wild (DW), periurban (P), pristine wild (PW) and urban (U).

### Entomological sampling

A total of 891 monitoring events (AAS x month) were performed, with 823 (92.5%) operating successfully, while the remaining 68 were either missing or had dried out. Of the successfully operating AAS x month, 19.6% (161) were positive for *Ae. albopictus*, and 39.9% (328) were positive for *Ae. aegypti*. In total, 1,614 *Ae. albopictus* specimens and 4,358 *Ae. aegypti* specimens were identified. The POI curve for *Ae. albopictus* exhibited a pronounced bimodal pattern, with the highest peak in January 2020 and a smaller peak in May 2019, while the lowest values were observed between June and October 2019, closely tracking precipitation patterns. For its part, the POI for *Ae. aegypti* dropped in autumn when monthly accumulated precipitation decreased but exhibited an oscillatory pattern during the winter dry season, then peaked in November 2019 and maintained a high plateau throughout the summer 2020 (Fig. 3A). As for temperature, lowest positive values for both POI coincide with low winter temperatures, especially Min_T. Some minor fluctuations were also detected in the NDVI and CI seasonal patterns (Fig. 3B).

**Figure 3.**
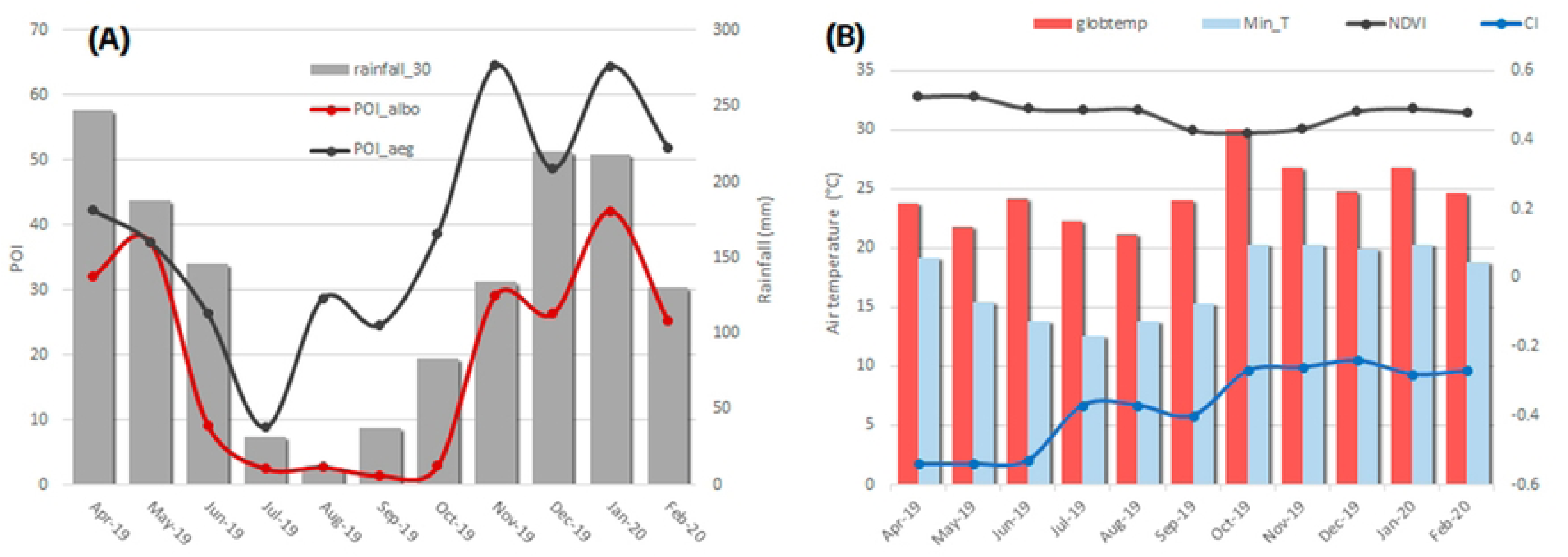
Monthly Positivity Ovitrap Index and Monthly variation spectral indices. (A) Monthly Positivity Ovitrap Index (POI) for *Ae. albopictus* and *Ae. aegypti*, along with monthly accumulated precipitation in bars. (B) Monthly variation of the spectral indices NDVI and CI; average air temperature (globtemp) and average minimal temperature (Min_T) in bars.

Regarding spatial heterogeneity, the POI for *Ae. albopictus* exhibited the highest values in periurban environments, particularly in the southern regions near the boundary of the Puerto Península Provincial Park (PPPP in Fig. 1) and showed moderate to low values in both urban and wild areas (Fig. 4A). Conversely, *Ae. aegypti* showed elevated values predominantly in urban environments, with a relatively uniform distribution, and moderate to low values in periurban areas. Notably, only one AAS recorded high values in the wild environment, located in a high pedestrian traffic zone (Fig. 4B).

**Figure 4.**
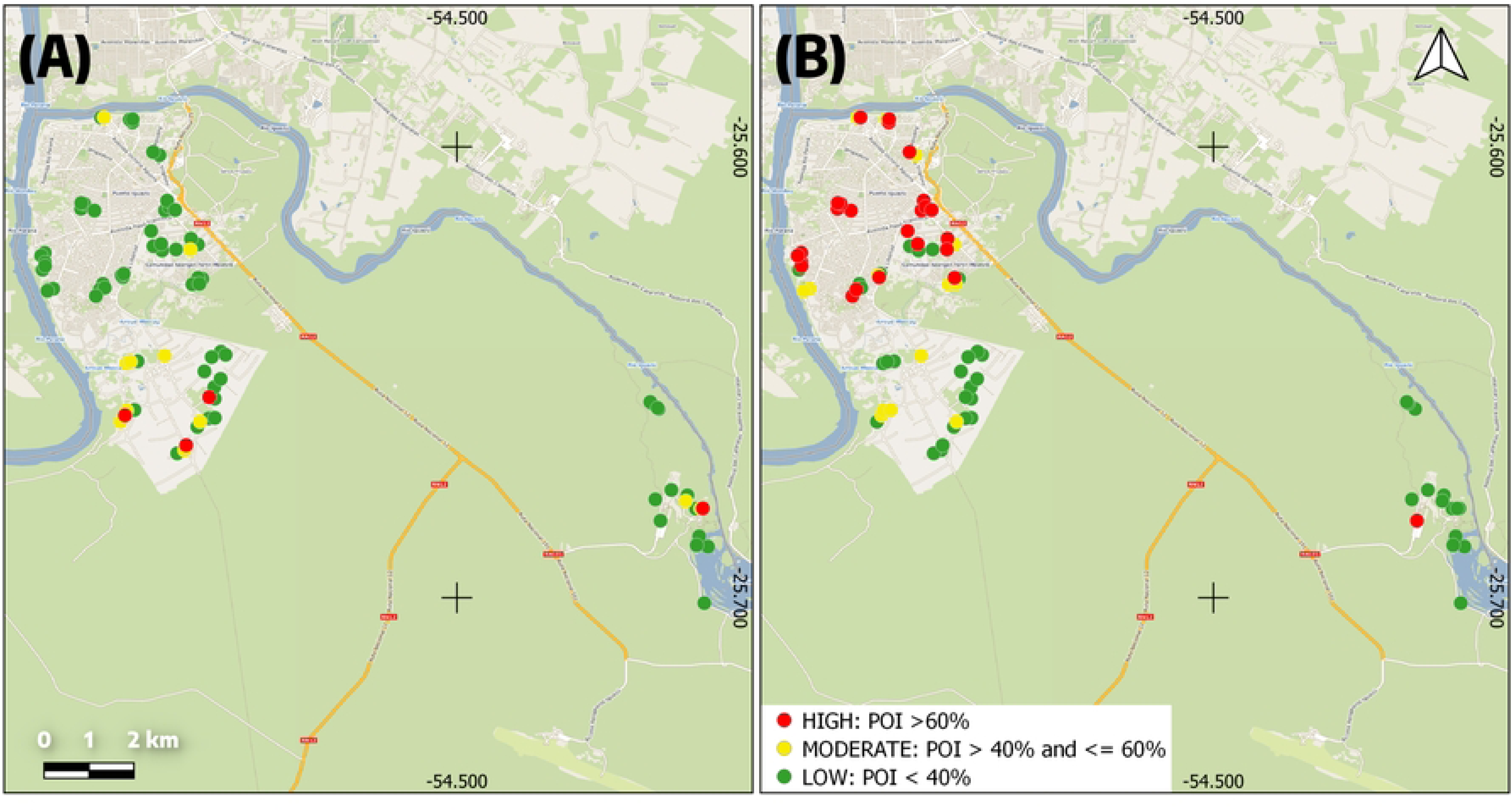
Location of the adult activity sensors (AAS). classified in three risk categories by their positivity ovitrap index (POI) values during the period April 2019 - February 2020 in Puerto Iguazú, Argentina, for *Ae. albopictus* (A) and *Ae. aegypti* (B).

### *Aedes* dynamics of environmental type

Two-thirds of the specimens of *Ae. albopictus* identified were collected in the periurban environment, followed by urban and disturbed wild at similar percentages (14-15%, Fig. 5A). The mean total abundance of *Ae. albopictus* per AAS was similar in disturbed wild (35.4) and periurban (31.7), and considerably lower in urban (7.8) and pristine wild (5.5). Highest POI values shifted from periurban in April-June to disturbed wild in July-December, and back to periurban in January-February (Fig. 5C).

**Figure 5.**
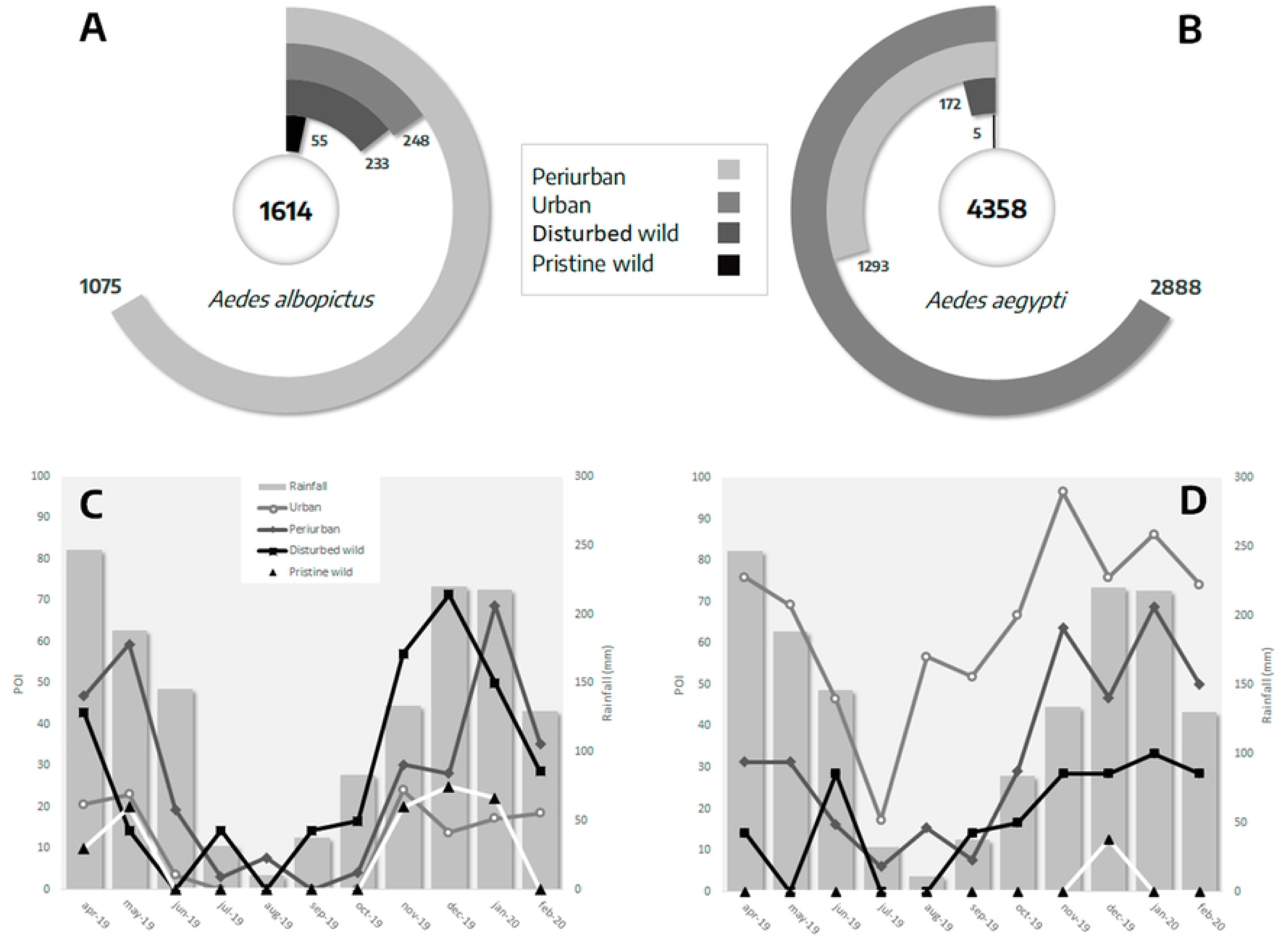
Total number of specimens of *Aedes albopictus*. (A) and *Ae. aegypti* (B) collected per environmental type, along with Positivity Ovitrap Index (POI) per month and environmental type overimposed on monthly cumulative precipitation (gray bars) for *Ae. albopictus* (C) and *Ae. aegypti* (D).

In contrast, the majority of *Ae. aegypti* specimens identified were collected in urban sites (Fig. 5B), with highest mean abundance per AAS (96.3) and highest POI values throughout all seasons, followed by periurban particularly during spring and summer (Fig. 5D) with a total mean abundance of 38.0. Values in disturbed and pristine wild environments (24.6 and 0.5, respectively) were lower compared to *Ae. albopictus*.

### Co-occurrence

The H0 was rejected for all environmental types except pristine wild, indicating an association between the presence of *Ae. albopictus* and *Ae. aegypti* in urban, periurban and disturbed wild environments. However, a significant association indicating coexistence (or avoidance), was only obtained in the urban (p < 0.001) and periurban (p = 0.027) environments. In both environmental types, the PCO was higher than the EO of *Ae. albopictus* (16% and 1% respectively in urban environments, 36% and 25% in periurban environments), indicating that the association between them is positive.

### Multivariate modeling of *Ae. albopictus* distribution

The correlation analysis among the explanatory variables (Supplementary data 2) showed that land cover classification variables, such as the extent of bare soil and high vegetation, exhibited high correlation values (ρ > 0.80). The latter (HgVeg) was chosen, as it better characterizes areas with minimal human intervention. Variables related to urbanization and the unemployment rate were also highly correlated, with impervious being selected due to its calculation from images closer to the sampling dates. Cumulative precipitation over 2 weeks and 30 days prior to the deactivation of the AAS were highly correlated, with the latter (rainfall_30) being selected. Between the average maximum and minimum temperature during the activation week of the AAS (Max_T and Min_T), the latter was retained, as it represents a more critical factor for larval development [56]. Finally, water and air temperatures showed ρ > 0.7; air temperature was chosen due to having more measurements (both during the activation and deactivation of the AAS).

Two alternative random structures were selected, both incorporating the season (N = 4) along with: (A) quadrants (N = 22); or (B) ID of the AAS (N = 81). In the univariate analyses, the two main variables are positively associated with the occurrence of *Ae. albopictus* were rainfall_30, the number of days that the AAS remained active (days) and Min_T. For the random structure (A), the categorical variable for road classification (Road_type) and the presence of *Ae. aegypti* (aegbin) were the most strongly associated, followed by urban land use class in a 150 m buffer around each AAS (Impervious), Land Surface Temperature (LST), air humidity (globhum), and environment (environment) to a lesser extent. For the random structure (B), along with rainfall_30 and days, environment and Impervious were equally significant, followed by LST, Road_type, distance to the nearest water body (Water_m), globhum, and the percentage of low vegetation in a 150 m buffer around each AAS (LowVeg) (see Supplementary Material 3).

Based on the specified random structures, three alternative models were attained: M1 and M2 with random structure (A), and M3 with random structure (B). The random effects accounted for 12%, 10%, and 17% of the variance in the occurrence of *Ae. albopictus*, respectively, while the fixed effects explained 47%, 50%, and 50%. The best-fitting model was M3, with a Kappa value of 0.63 (Table 2). In it, monthly precipitation exhibited a positive quadratic association that led to an increase in the probability of occurrence of *Ae. albopictus* from approximately 20% with 10.5 mm of monthly cumulative precipitation values to 54% at peak values of 64.5 mm, followed by a decline at higher precipitation values. Regarding temperature, LST showed a nonlinear inverse association, meaning that as temperature increases, the probability of presence of *Ae. albopictus* decreases, albeit with minimal influence, reaching maximum probability values (0.4%) at 16.12°C. The variable days was positively associated, indicating a higher probability of occurrence of *Ae. albopictus* as the number of exposure days of the AAS increased. LowVeg was selected as a quadratic association, with a growth area ranging from 33.8% when LowVeg is 0, to a maximum probability value of 35.5% at 5% LowVeg coverage and decreasing again to a 0% probability of occurrence of *Ae. albopictus* at coverage values around 60%. Impervious exhibited a nonlinear inverse relationship with the occurrence of *Ae. albopictus*, with maximum probability values of 34% when Impervious is 0% and gradually decreasing as impervious increases. The presence of *Ae. aegypti* was associated with the probability of occurrence of *Ae. albopictus* in the random structure models (A), taking probability values of M1=43.3% and M2=30%. Lastly, for these two models, salinity (sal) was also selected, with probabilities ranging from ≅ 23% when sal=0.01 ppt to M1=93% and M2=83.4% when sal=1.01 ppt in both cases (Fig. 6).

**Figure 6.**
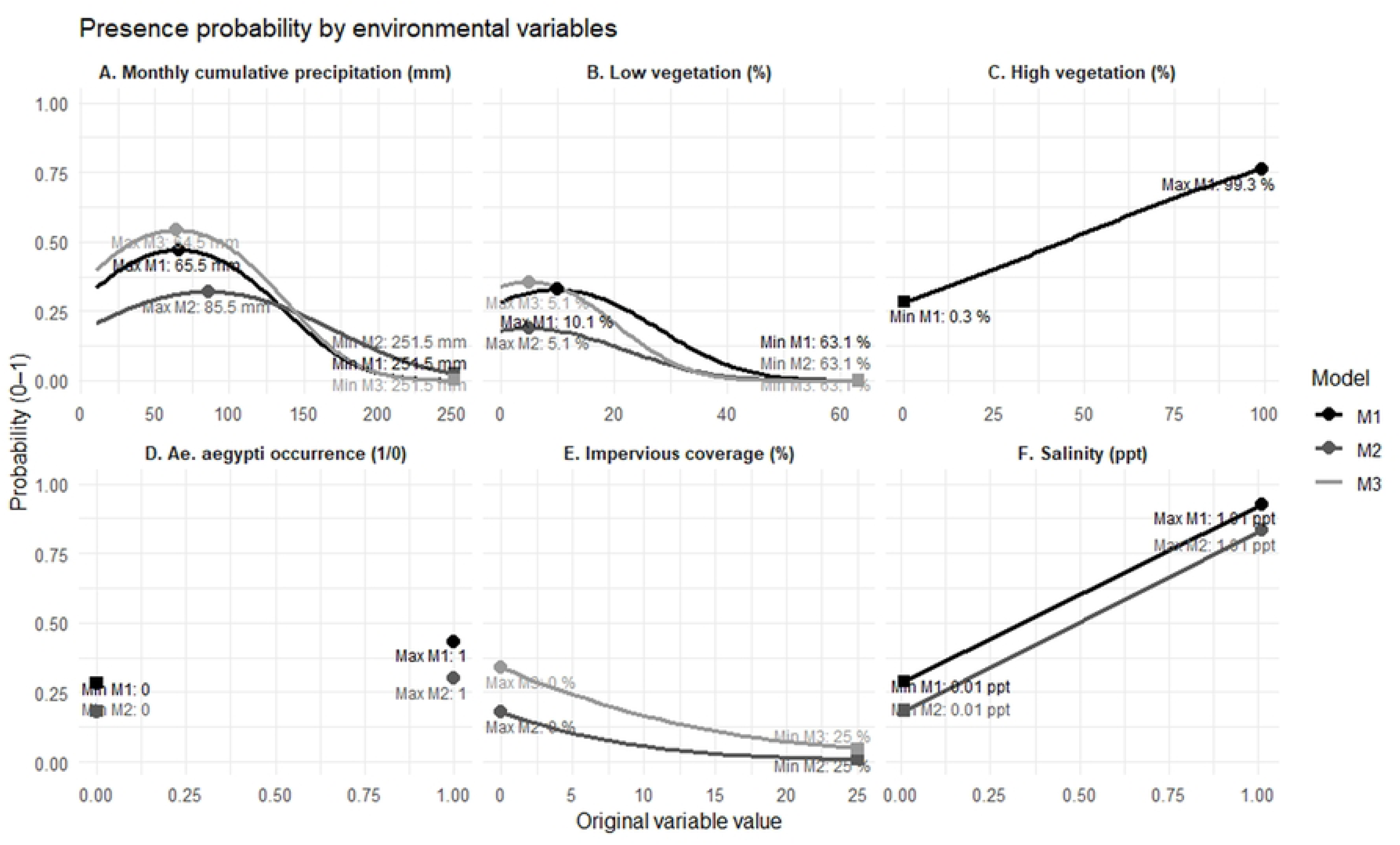
Probability of presence of *Aedes albopictus* as a function of environmental variables, according to the best-fitted generalized linear mixed models (M1, M2, and M3). Panels correspond to: (A) Monthly cumulative precipitation (mm), (B) Low vegetation (%), (C) High vegetation (%), (D) *Ae. aegypti* occurrence (1/0), (E) Impervious coverage (%), and (F) Salinity (ppt). Values of the explanatory variables that predict the maximum and minimum probability of occurrence of *Ae. albopictus* are indicated for each model.

**Table 2.**
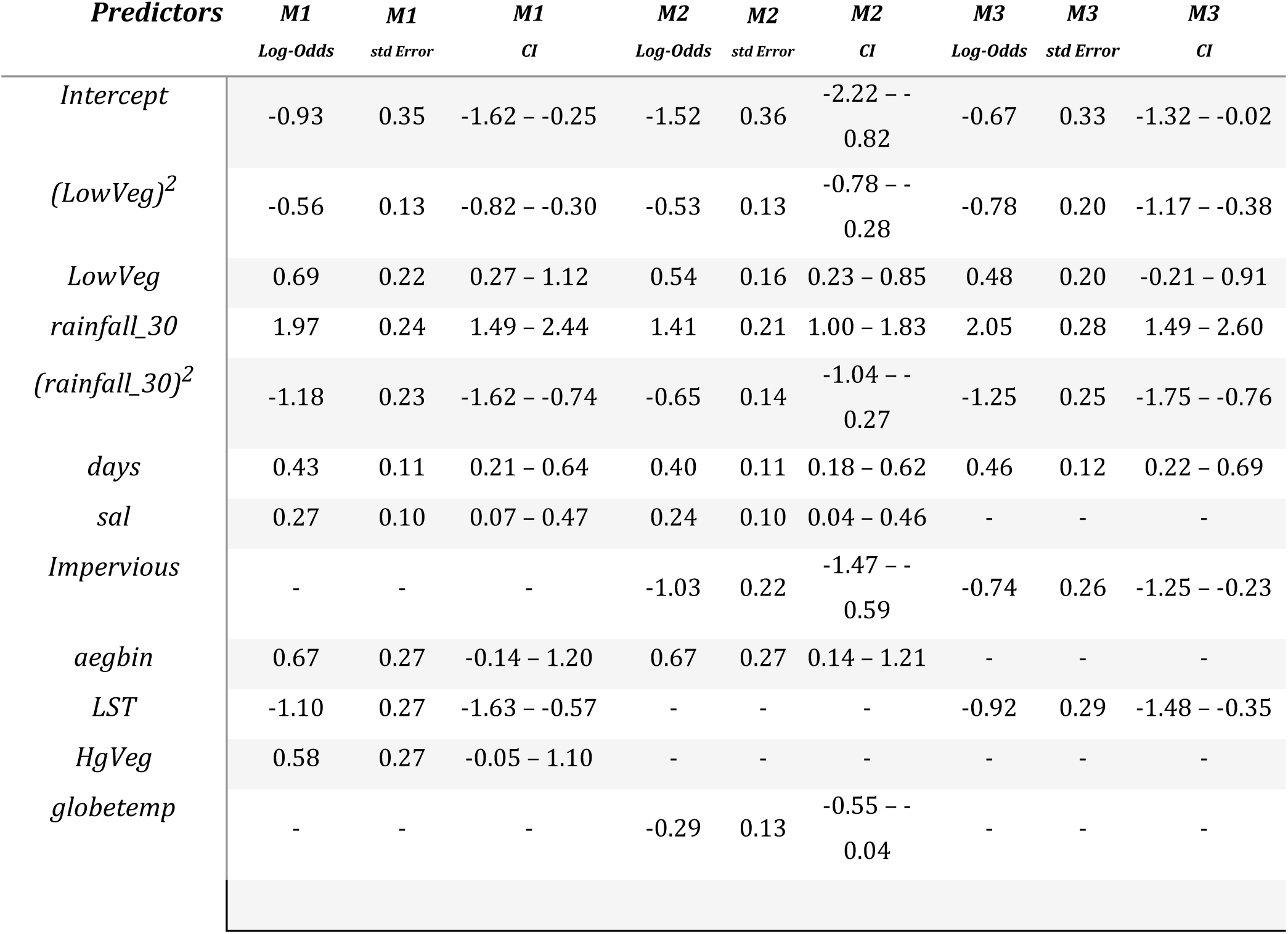
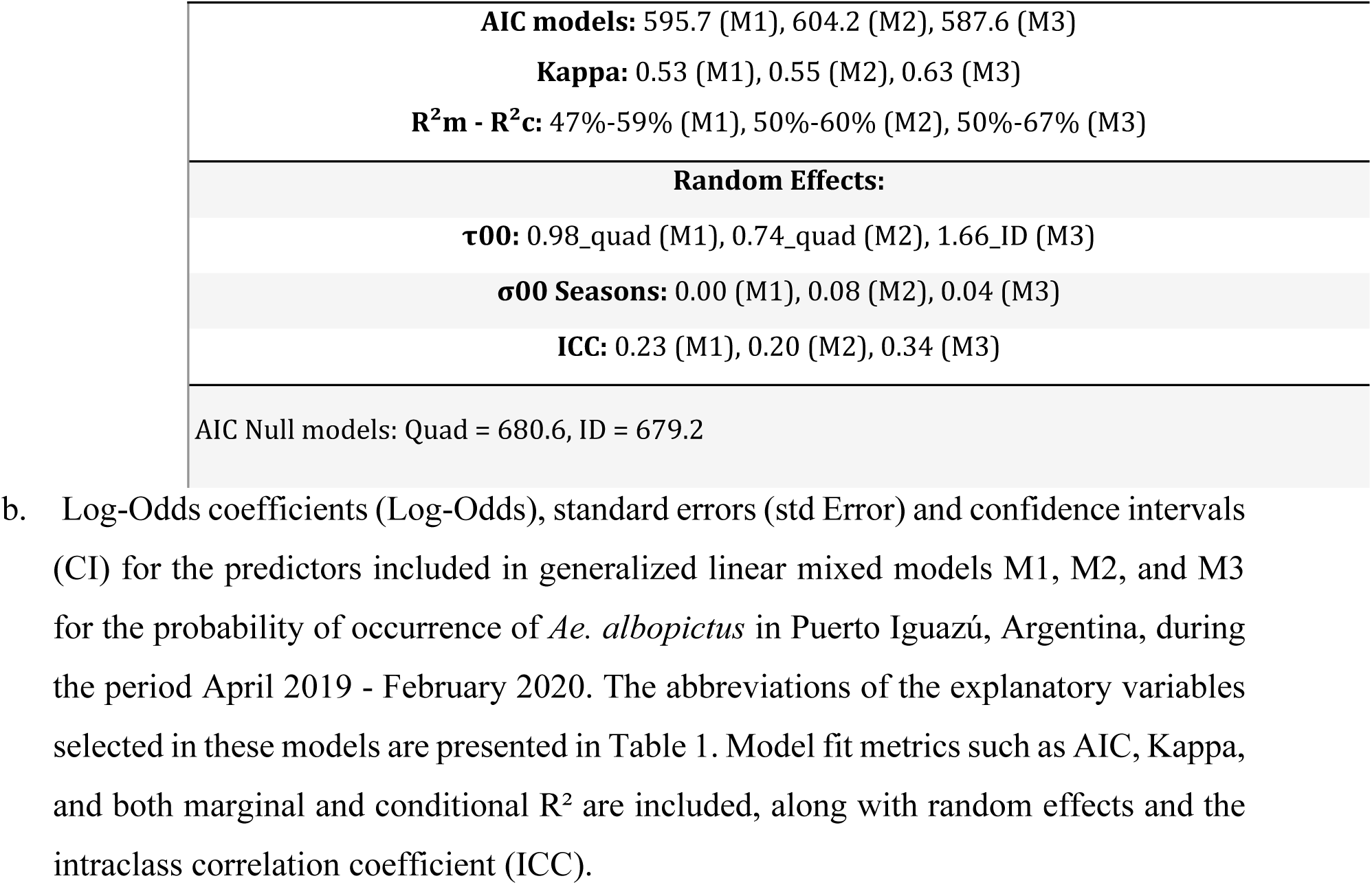
Log-Odds coefficients (Log-Odds), standard errors (std Error) and confidence intervals (CI).

| *Predictors* | *M1*<br>*Log-Odds* | *M1*<br>*std Error* | *M1*<br>*CI* | *M2*<br>*Log-Odds* | *M2*<br>*std Error* | *M2*<br>*CI* | *M3*<br>*Log-Odds* | *M3*<br>*std Error* | *M3*<br>*CI* |
| --- | --- | --- | --- | --- | --- | --- | --- | --- | --- |
| *Intercept* | -0.93 | 0.35 | -1.62 – -0.25 | -1.52 | 0.36 | -2.22 – -0.82 | -0.67 | 0.33 | -1.32 – -0.02 |
| *(LowVeg)<sup>2</sup>* | -0.56 | 0.13 | -0.82 – -0.30 | -0.53 | 0.13 | -0.78 – -0.28 | -0.78 | 0.20 | -1.17 – -0.38 |
| *LowVeg* | 0.69 | 0.22 | 0.27 – 1.12 | 0.54 | 0.16 | 0.23 – 0.85 | 0.48 | 0.20 | -0.21 – 0.91 |
| *rainfall_30* | 1.97 | 0.24 | 1.49 – 2.44 | 1.41 | 0.21 | 1.00 – 1.83 | 2.05 | 0.28 | 1.49 – 2.60 |
| *(rainfall_30)<sup>2</sup>* | -1.18 | 0.23 | -1.62 – -0.74 | -0.65 | 0.14 | -1.04 – -0.27 | -1.25 | 0.25 | -1.75 – -0.76 |
| *days* | 0.43 | 0.11 | 0.21 – 0.64 | 0.40 | 0.11 | 0.18 – 0.62 | 0.46 | 0.12 | 0.22 – 0.69 |
| *sal* | 0.27 | 0.10 | 0.07 – 0.47 | 0.24 | 0.10 | 0.04 – 0.46 | - | - | - |
| *Impervious* | - | - | - | -1.03 | 0.22 | -1.47 – -0.59 | -0.74 | 0.26 | -1.25 – -0.23 |
| *aegbin* | 0.67 | 0.27 | -0.14 – 1.20 | 0.67 | 0.27 | 0.14 – 1.21 | - | - | - |
| *LST* | -1.10 | 0.27 | -1.63 – -0.57 | - | - | - | -0.92 | 0.29 | -1.48 – -0.35 |
| *HgVeg* | 0.58 | 0.27 | -0.05 – 1.10 | - | - | - | - | - | - |
| *globetemp* | - | - | - | -0.29 | 0.13 | -0.55 – -0.04 | - | - | - |

|  |
| --- |
| **AIC models:** 595.7 (M1), 604.2 (M2), 587.6 (M3) |
| **Kappa:** 0.53 (M1), 0.55 (M2), 0.63 (M3) |
| **R<sup>2</sup>m - R<sup>2</sup>c:** 47%-59% (M1), 50%-60% (M2), 50%-67% (M3) |
| **Random Effects:** |
| **τ00:** 0.98_quad (M1), 0.74_quad (M2), 1.66_ID (M3) |
| **σ00 Seasons:** 0.00 (M1), 0.08 (M2), 0.04 (M3) |
| **ICC:** 0.23 (M1), 0.20 (M2), 0.34 (M3) |
| AIC Null models: Quad = 680.6, ID = 679.2 |

The geostatistical projections of model M3 are shown in Figure 7. The IDW interpolation method was employed during the winter season for both the INP and non-INP groups due to the absence of significant spatial autocorrelation, as evidenced by Moran’s I values ranging from −0.05 to 0.04 (p ≥ 0.06). In autumn, Kriging was applied for the non-INP group, where short-range spatial dependence was evident (∼0.9 km, Moran’s I = 0.20, p = 1.0 × 10⁻⁶; ∼2.6 km, Moran’s I = 0.06, p = 0.009), while IDW was maintained for INP (p > 0.3). In the spring, Kriging was employed for both groups due to the presence of significant positive correlations at short distances (INP: ∼1.3 km, Moran’s I = 0.10, p = 0.0025; non-INP: ∼0.9–2.6 km, Moran’s I = 0.12–0.10, p < 0.001). During the summer season, Kriging was employed for the non-INP group, given the pronounced clustering observed (∼0.9 km, I = 0.27, p = 1.1 × 10^-10^).

**Figure 7.**
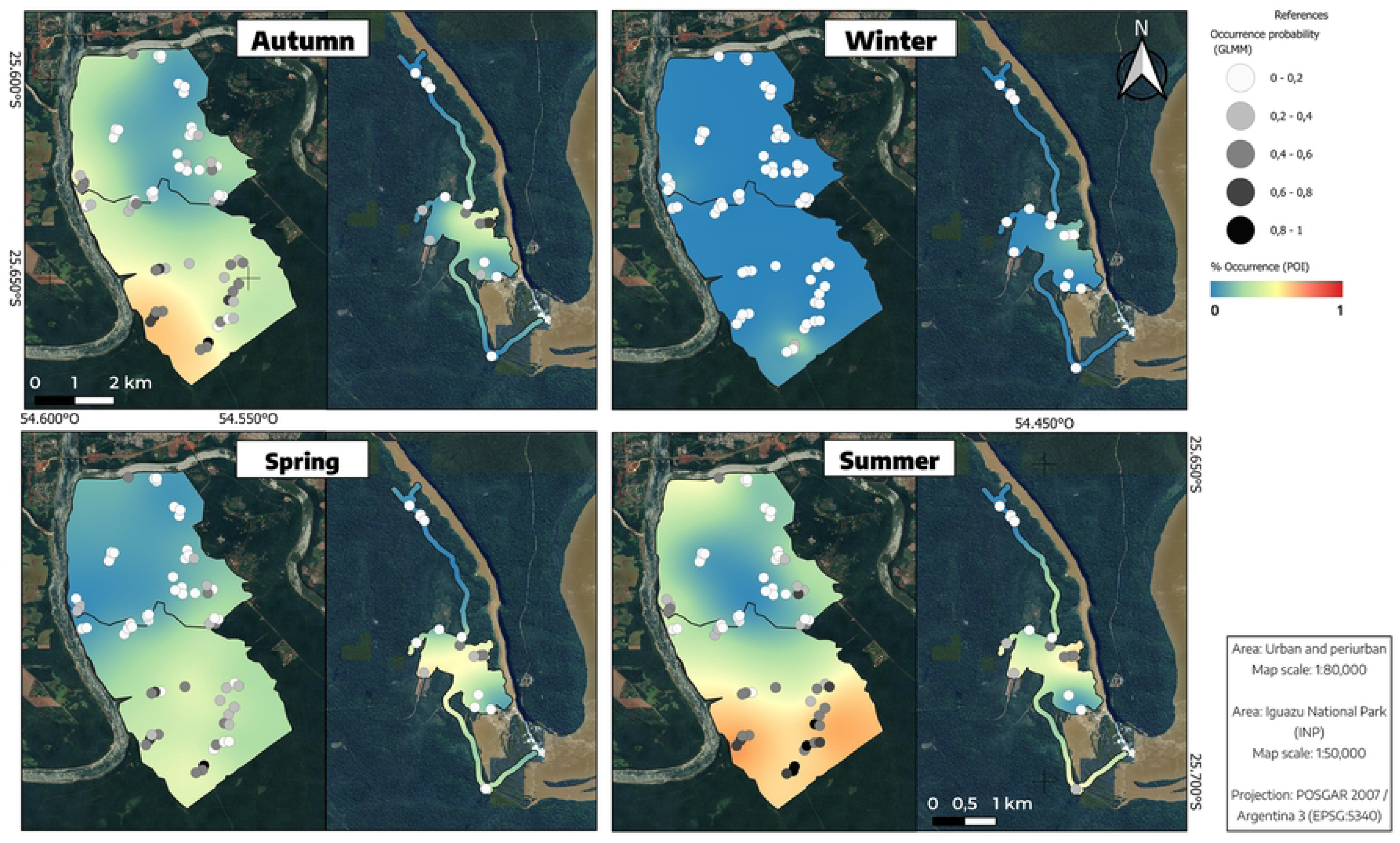
Spatial interpolations using Kriging and IDW of the probability of occurrence of *Aedes albopictus*, based on POI values and predictions from model M3. The points represent the locations of the AAS, with their predicted probabilities (0–1) shown in grayscale. The interpolated probability of occurrence is displayed with a color gradient ranging from blue tones (low probability, close to 0) to red tones (high probability, close to 1).

Conversely, IDW was maintained for INP, as the calculation of a correlogram was precluded by data limitations. Autumn and spring exhibit similar probabilities of occurrence, although the trend suggests that the species remains more prevalent after summer, whereas its distribution after winter becomes spatially more restricted. At the spatial level, a clear zoning pattern is observed, with a major hotspot located in the peri-urban environment near the PPPP and PHDNR reserves (see Fig. 1). This area is characterized by lower housing density and infrastructure, while patches of wild vegetation are preserved and intermixed with intensive subsistence agriculture. During periods of higher probabilities, the borders of the urban area “light up”, but the species is virtually absent in the central urban core, where urbanization is more consolidated. The disturbed wild environment, particularly the public-use sector of INP, shows a localized hotspot in autumn that persists slightly into winter and re-emerges in spring, spreading more evenly and continuing into summer. In contrast, in the pristine wild environment, almost no probability of occurrence is observed throughout the seasons, except for a slight increase in the southern part of the area closest to the circuits with the highest human traffic in the INP during summer. Projected predictions for models M1 and M2 were consistent with those of M3 (data not shown).

## Discussion

Findings in sub-tropical Argentina provide robust evidence for a spatially stratified distribution of *Ae. albopictus*, with the periurban environment acting as the most favorable setting for its proliferation. This pattern aligns with previous work documenting increased vector presence in ecotones and buffer zones [57; 26]. Transitional environments, where anthropogenic landscapes interlace with remnants of native forest, create pronounced edge effects that appear to facilitate vector presence—an observation also documented for *Haemagogus* spp., a sylvatic vector of yellow fever, in similar contexts [58].

Consistent with the observations of Lazaric et al. [59], the area labelled herein as disturbed wild, located within public use zones of the INP, is associated with features typical of periurban environments and also harbored *Ae. albopictus* throughout the year. In this context, vector presence appears to be associated with anthropogenic interventions that lead to land-use changes affecting natural environments. In contrast, the spatial stratification observed for *Ae. aegypti* demonstrates a clear dominance of the species in highly urbanized settings, in coincidence with the literature [19] and supported by the co-occurrence results presented here, which revealed a positive association between both species, with *Ae. aegypti* dominating in urban environments. Regarding temporal distribution, our study revealed *Ae. aegypti* oviposition throughout the year in both urban and periurban environments, could serve as a strong predictor of dengue endemicity, however, endemic dengue requires the concurrence of additional factors, such as persistent virus circulation, adequate susceptible host density, and suitable climatic conditions [60]. This finding underscores the need to intensify studies in other localities to compare and validate these results.

Building on Espinosa et al. [57], we observed that, as urbanization expanded and intensified over space and time within the study area, *Ae. albopictus* progressively shifted toward more recently urbanized zones with lower levels of imperviousness. Consistent with Reiskind et al. [61], the species exhibited greater stability at the edges of densely vegetated areas. Therefore, land-use changes can be considered a risk factor, and their identification and monitoring should be regarded as an early warning indicator of potential colonization by *Ae. albopictus*. This highlights the critical role of natural environments in maintaining ecosystem functions with direct implications for public health, as land-use change and habitat fragmentation have been identified as contributing factors in the emergence and transmission of zoonotic diseases [30; 62].

When analyzing our results in conjunction with previous studies in the region, we observe heterogeneity in vector distribution as a function of urbanization. Lizuain et al. [13] assessed the occurrence of *Ae. albopictus* in two localities with contrasting infrastructure and urban development: Eldorado (26°24′00″S; 54°38′00″W), a large city with high urban density, and Colonia Aurora (27°28′29″S; 54°31′28″W), a smaller town with a rural profile and greater connectivity to natural areas. In Eldorado, *Ae. albopictus* was reported with low relative abundance, whereas in Colonia Aurora, it showed the highest relative abundance of *Ae. albopictus* recorded in Argentina to date in co-dominance with *Ae. aegypti*. Accordingly, the abundance of *Ae. albopictus* across these localities, along with the areas examined in the present study, can be represented by a bell-shaped pattern, as illustrated in Figure 8: low to moderate presence in Eldorado and the urban area of Puerto Iguazú, gradually increasing in the periurban of Puerto Iguazú, peaking in Colonia Aurora, then declining in the disturbed wild areas of the INP, and reaching minimal levels in the pristine wild zones of the INP [13].

**Figure 8.**
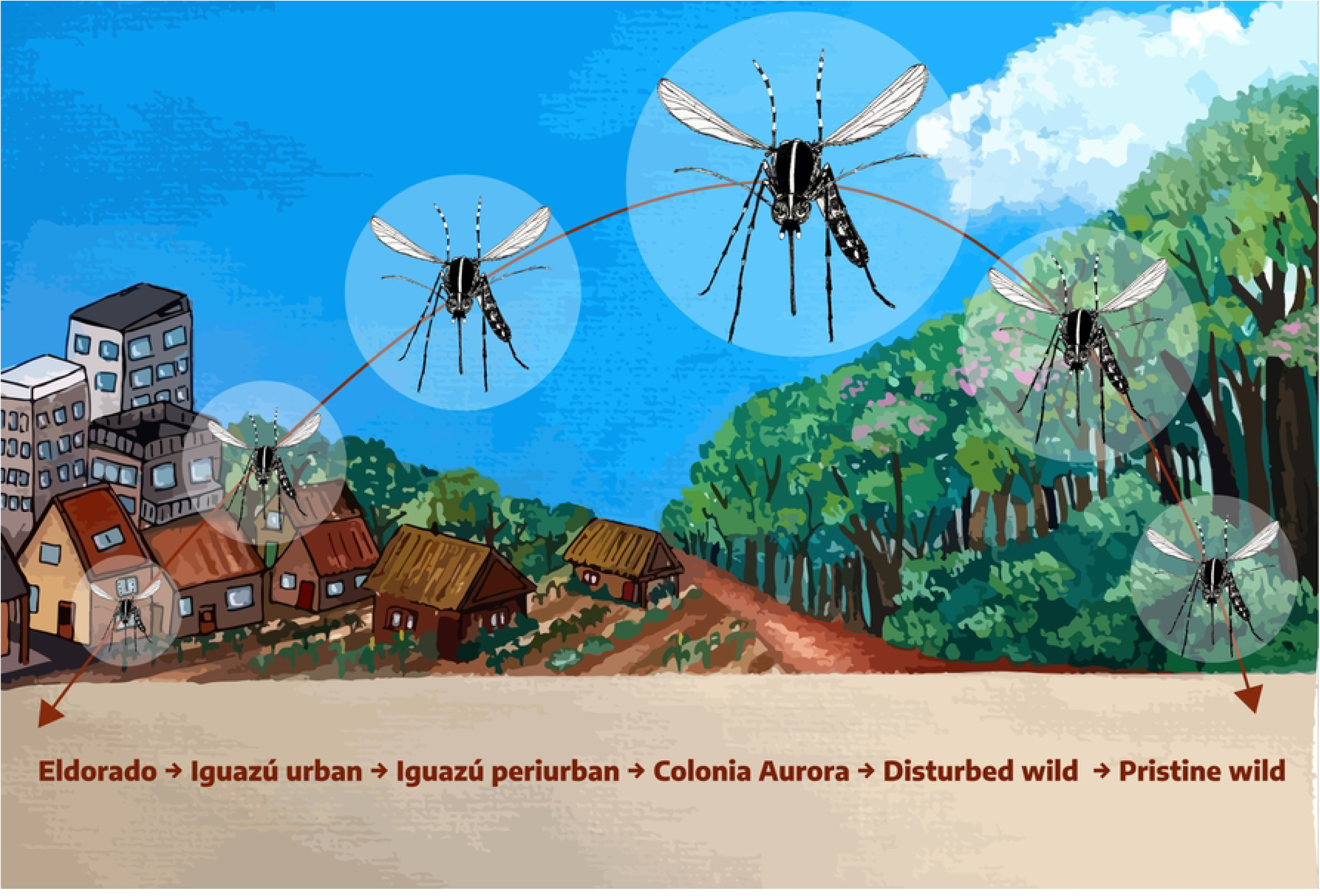
Urbanization gradient of localities in Misiones, Argentina, with reported presence of *Aedes albopictus*. The sequence from higher to lower urbanization includes: Eldorado, Iguazú urban, Iguazú peri-urban, Colonia Aurora, disturbed wild at Iguazu National Park, and pristine wild at Iguazu National Park. The species shows moderate to low presence in the most urbanized areas, reaching its maximum occurrence in Colonia Aurora, and decreasing towards native forest with minimal human intervention.

Seasonal dynamics of *Ae. albopictus* was primarily driven by precipitation, as previously reported [19; 63], presumably due to water availability acting as a limiting factor in peridomestic environments [28], while minimum temperature thresholds had a secondary influence. At the microhabitat scale, the physicochemical properties of water within containers were also associated with *Ae. albopictus* presence. Higher salinity was positively associated, likely reflecting the accumulation of decomposing organic matter. Organic detritus releases dissolved ions—such as nitrates, phosphates, and humic substances—that enrich aquatic habitats and enhance larval development [64]. This process was especially evident in ecotonal areas, where containers are frequently surrounded by vegetation, contrasting with urban environments dominated by clean water in artificial containers. Future studies should address this interaction between physicochemical and environmental factors, which may help explain why *Ae. albopictus* tends to establish more strongly in transitional areas with anthropogenic influence and surrounding vegetation, compared to densely urbanized settings or pristine forest environments. Importantly, none of the oviposition sensors deployed during the study were found to contain *Aedes* larvae or pupae, as they were deactivated before egg hatching.

Upon analysis of the spatial autocorrelation results, it was evident that this phenomenon was manifested most prominently during the spring and summer months. During the autumn season, this effect was observed to a lesser extent, however, this observation pertained exclusively to the non-INP group. The winter season, conversely, exhibited an absence of any discernible autocorrelation. In the non-INP group, two significant distance classes were detected (∼ 0.9 km and ∼ 2.6 km), suggesting the existence of two clustering patterns, possibly associated with the combination of urban and periurban environments within the same group. These results are consistent with the seasonal patterns of probability of occurrence of *Ae. albopictus*, which peaks in summer and decreases almost completely in winter.

This study contributes to the understanding of the environmental factors influencing the spatiotemporal distribution of *Ae. albopictus*, in order to support risk stratification for the transmission of zoonotic diseases in the tri-border area of Argentina, Brazil, and Paraguay.

Based mostly on remote sensing data, we defined an environmental stratification of the study area into four sectors of epidemiological interest. The ability to stratify environments by identifying variables that characterize ecological or demographic areas of interest, and that show significant correlation with the spatiotemporal distribution of *Ae. albopictus*, represents a valuable monitoring tool for entomological risk stratification and for guiding zoonotic disease prevention efforts.

A strong association was found between the spatiotemporal distribution of *Ae. albopictus* and land-use changes resulting from unplanned urban development. These landscapes represent high-risk scenarios for the spread of zoonotic diseases. Colonization of the vector is postulated to be driven by a delicate balance within the landscape, wherein moderate human intervention provides access to a diversity of larval habitats (both natural and artificial), while maintaining sufficient connectivity to native forest patches or dense and heterogeneous vegetation that offer blood sources and resting sites. We propose that the colonization of new environments by *Ae. albopictus* is significantly associated with recent anthropogenic disturbance in wild areas, and that the species exhibits metapopulation dynamics—where periurban zones function as source habitats, while both anthropized and pristine areas act as sinks.

Thus, the risks associated with zoonotic disease transmission are not only linked to socioeconomic conditions and access to basic public services, but also to local socio-environmental factors—such as the absence of political-administrative tools for environmental land-use planning—and regional variables related to climate dynamics. We hypothesize that territorial planning strategies could mitigate health risks. In line with the “urban One Health” paradigm proposed by Ellwanger et al. [65], promoting sustainable urbanization with adequate infrastructure may not only limit mosquito proliferation but also improve broader population health outcomes.

## Conclusions

In conclusion, our findings support the that the establishment of *Ae. albopictus* requires specific environmental conditions, particularly in areas undergoing frontier expansion without proper environmental planning. Therefore, we recommend adopting an integrated One Health approach: multiscale approach that combines ecological surveillance, remote sensing, and land-use planning offers a powerful framework for proactive vector control. Such strategies are particularly relevant in biodiverse and ecologically sensitive regions like the tri-border area of Argentina, Brazil, and Paraguay, where zoonotic disease risks intersect with rapid territorial transformation.

## Acknowledgments

We gratefully acknowledge Dr. Lucía Maffey and Lic. Graciela Minardi for their insightful suggestions, language corrections, and overall contributions to improving the manuscript. We also extend our thanks to IMIBIO, the Province of Misiones, and the APN —particularly Iguazú National Park and its rangers—for granting research permits and for their continuous support and collaboration throughout the study.

## Data availability statement

The data supporting the findings of this study are available from the corresponding author upon reasonable request.

## Author contributions

J.A. Siches led the conceptualization of the study and was responsible for data collection, formal analysis, and visualization. She developed the original draft of the manuscript and contributed to the development of the methodology, project administration, validation, and the processing of satellite imagery.

M.V. Micieli contributed to project administration, supervision, manuscript review and editing, and the provision of resources.

P.E. Berrozpe participated conceptualization and data collection, investigation, methodological design, validation, supervision, and manuscript review and editing.

R. Iglesias assisted with data curation, processing of satellite imagery, and visualization.

J.J. García contributed to conceptualization, resource provision, project administration, and supervision.

M.V. Cardo contributed to conceptualization, data curation, formal analysis, and mathematical model development, and participated in manuscript review and editing.

All authors contributed to the interpretation of results and approved the final version of the manuscript.

